# Organism-wide mapping of MHC class I and II expression in mouse lemur cells and tissues

**DOI:** 10.1101/2022.02.28.482372

**Authors:** Lisbeth A. Guethlein, Camille Ezran, Shixuan Liu, Mark A. Krasnow, Peter Parham

**Affiliations:** Department of Structural Biology, Stanford University School of Medicine, Stanford, CA, USA; Department of Microbiology and Immunology, Stanford University School of Medicine, Stanford, CA, USA; Department of Biochemistry, Stanford University School of Medicine, Stanford, CA, USA; Howard Hughes Medical Institute, USA; Department of Chemical and Systems Biology, Stanford University School of Medicine, Stanford, CA, USA

## Abstract

The major histocompatibility complex (MHC) class I and II glycoproteins have been associated with numerous disease phenotypes, mechanisms and outcomes. These associations can be due to allotypic polymorphism or to altered levels of allotype expression. Although well studied in a range of cell types and microenvironments, no study has encompassed all cell-types present in an individual mouse lemur. In this study the droplet-based single cell RNA sequence data from the mouse lemur cell atlas, Tabula Microcebus, were used to examine the patterns of MHC class I and II expression. The cell atlas comprises data obtained from 27 organs from four mouse lemurs, enabling comparison of expression pattern between both cell types and the individual lemurs. Patterns of gene expression showed a good concordance among the four mouse lemurs. Three primary patterns of expression were identified and associated with different cell-types. Mapping MHC expression onto existing cell trajectories of cell development and spatial gradients revealed fine scale differences in expression level and pattern in single tissues. The bioinformatic pipeline developed here is applicable to other cell atlas projects.

## INTRODUCTION

In the genomes of vertebrate species the major histocompatibility complex (MHC) is a region densely populated with genes that contribute to innate and adaptive immunity. Defining the MHC are highly polymorphic MHC class I and II glycoproteins that bind peptides derived from viruses and other intracellular pathogens, that can then activate the immune response of natural killer (NK) cells and T cells^1,2^. Differences between MHC class I and II allotypes often involve multiple amino acid substitutions, whereas most polymorphisms of immune system proteins involve a single substitution. Consequently, many more diseases are associated with MHC class I and II gene polymorphisms, than with polymorphism in any other immune system genes^3^.

The MHC class I and II cell-surface glycoproteins present peptides to receptors of the immune system and through these interactions modulate a variety of downstream responses. MHC class I molecules also function as ligands for receptors on Natural Killer (NK) cells, through generation of the ‘missing self’ response. This response of innate immunity detects changes in cell surface expression of MHC class I, causing cells with altered or reduced cell-surface expression to be eliminated. Both MHC class I and II present peptide antigens to T-cells, thereby serving as ligands for the αβT cell receptors that activate adaptive immune responses.

The *MHC class I* and *II* genes are some of the most polymorphic genes in primates. The mouse lemur is no exception as is seen from the diversity of *Mimu-DRB* alleles that emerged from the study of wild lemur populations^4–7^. Mouse lemurs, one of the world’s smallest primates, are strepsirrhine primates. Their lineage diverged from the human lineage 64 mya^8^. The pioneer study of mouse lemur *MHC class I* defined four distinctive *MHC class I* sequences: *Mimu-W01, -W02, -W03* and *-W04*^9^. The extent to which these sequences represented either different genes or different alleles was uncertain. Averdam et al^10,11^ sequenced several BACs containing *MHC class I* and/or *class II* genes. Within the region of the mouse lemur *MHC* that is syntenic to the *class I* region of other primates, the *class I* genes are all predicted to be nonfunctional. In contrast, those *MHC class I* genes that are predicted to encode functional MHC class I proteins cluster on a different chromosome from that encoding pseudogenes. The paired genes, encoding mouse lemur *MHC class II* alpha and beta chains, are in a region syntenic to *MHC class II* region of human chromosome 6. Complicating the measure of MHC expression from short read sequencing data is the extensive polymorphism of individual genes and the high degree of homology among the genes. Although knowledge of mouse lemur *MHC class I* and *II* is limited, the general rule for other primate species is that *MHC class I* variation and polymorphism exceed that of *MHC class II*^12^.

Expression of MHC class I and II genes has been examined in a variety of tissues, cell types and cell environments^13^. By contrast, no previous study has comprehensively examined *MHC class I/II* expression across multiple organs and individuals at single cell resolution. To do this, we developed bioinformatic methods that determine MHC class I and II expression levels from 10x droplet-based scRNAseq data. These methods were applied to the Tabula Microcebus cell atlas (https://tabula-microcebus.ds.czbiohub.org/about)^14^, which comprises single cell transcriptomes from 27 organs and tissues of four individual mouse lemurs (*Microcebus murinus*). We analyzed 215,524 cells, representing 267 cell annotations and 739 molecular cell types. The Tabula Microcebus cell atlas enabled us to compare MHC class I and II gene expression across the tissues of one individual, as well as expression of the same MHC class I gene in four individuals.

In the course of analysing *MHC class I* and *II* gene expression for the mouse lemur cell atlas, we compared the alignments of Averdam et al^10,11^ to the most recent genome assembly^15^ (Mmur_3.0, https://www.ncbi.nlm.nih.gov/assembly/GCF_000165445.2). In the genome assembly, analysis of the *MHC* regions was complicated, because they were constructed from sequences derived from 4 individuals^15^. The NCBI annotation of the *MHC* region contained several genes and gene fragments with assigned homology to various *MHC* genes of different species. We first compared the BAC and genome sequences to assess potential allelic and structural variation in the *MHC* regions. This facilitated development of the bioinformatic pipeline that we used to determine gene content and polymorphism for the *MHC* genes of the four mouse lemurs used to construct Tabula Microcebus. These results were compared to the two genome sequence references to determine the accuracy of their assembly and annotation. The new techniques we developed should have broad application in studying *MHC class I* and *II* expression in the mouse lemur, as well as in other cell atlas projects.

## METHODS

### Reference dataset

*MHC class I* and *class II* gene sequences were obtained from Genbank. They comprise known expressed *class I* sequences^9^, sequences extracted from BACs that include full length genomic sequences^10,11^ and sequences annotated by NCBI (Refseq Annotation Release 101) in the mouse lemur genome assembly^15^ (Mmur_3.0). These were used to analyse genomic organization and allelic variation. Accession numbers of the sequences are in Supplementary Table I.

### Genomic organization

Segments of the mouse lemur *MHC* were sequenced previously from a series of BACs^10,11^. We compared these sequences to the corresponding regions of Mmur_3.0, by extracting the region from Mmur_3.0 and analysis using dotmatcher on the Galaxy server (https://usegalaxy.org/) with windowsize = 100 and threshold = 95. We also compared larger regions for marker order and synteny with the human genome (GRCh38).

All predicted mouse lemur *MHC class I* genes annotated by NCBI in Mmur_3.0, were extracted and aligned to the genes described previously^9,11^. Phylogenetic analysis was performed using this alignment and Mega 7^16^ (https://www.megasoftware.net). Neighbor-joining trees were constructed using 1000 bootstrap replicates, pairwise deletion and the Tamura-Nei substitution model. As two of the predicted sequences (XM_020282712 and XM_020282713) do not overlap, separate trees were constructed for each of these sequences and the results compared and combined. Predicted *MHC class II* gene sequences were extracted from Mmur_3.0 and aligned to the previously described mouse lemur *class II* genes and representative human *MHC class II* genes.

### MHC class I and class II sequence extraction from 10x scRNAseq data

Bowtie2^17^ was used to extract all reads mapping to *MHC class I* or *II* genes in the 10x scRNAseq datasets using --very-sensitive settings (http://bowtie-bio.sourceforge.net/bowtie2/manual.shtml). The reference used to build the index was a compilation of the known or predicted cDNA sequences for all mouse lemur *MHC class I* and *II* genes. The extracted reads for each individual were combined and then mapped to a reference comprising one example of each of the mouse lemur *MHC class I* (11 total) and *class II* genes (9 total). The resulting alignments were inspected manually using the Integrated Genomics Viewer^18,19^ (https://software.broadinstitute.org/software/igv) to determine allelic variants, the distribution of mapping quality, and the possibility of variation in the presence/absence, and/or copy number, of any of the genes.

Comparison of the lemur MHC *class I* sequences (Supplemental Figure 1) shows instances of intergenic recombination. Due to the length of the individual sequencing reads and the uneven distribution of reads coming from the 10x pipeline, unambiguous assignment of reads across the entire length of the *MHC class I* genes was not possible. In our expression analysis only 600-650 bp at the 3’ end could be assigned with confidence to a particular class I gene. This region covered over 99% of reads obtained in the 10x screen. Using the results from the global analysis, reference sequences were constructed for each individual lemur. Genes not present in an individual were omitted, so as to avoid mismapping of homologous reads to the reference sequence of absent genes. The reference sequences were analysed using a bioinformatic pipeline to determine the number of unique reads per cell for each gene in the reference.

The pipeline was applied separately to each individual-tissue combination. In the first step, bowtie2 (with --very-sensitive) was used to extract all reads mapping to the reference. Samtools^20^ was then used to sort the reads into separate files according to the gene to which they mapped. For some *class I* genes there were reads that could not confidently be assigned to just one gene, because of the high sequence homology between the various *class I* genes. Taking a conservative approach, we chose to remove those reads from the analysis. This was achieved following separation of the gene-specific files. Samtools was used to extract those reads with a mapping quality (MAPQ) score above a threshold determined in the course of the global sequence comparison. For each of the final read files, samtools was used to extract a list of the reads in each file. Seqtk (https://github.com/lh3/seqtk) was then used to extract the barcode and UMI for each of the reads in the readlist file. The duplicate barcode-UMI (unique molecular identifier) combinations were removed and the number of UMI per barcode was calculated. Final read counts were merged with the annotation file for the complete cell atlas.

In this paper, read counts have been normalized and are displayed as read counts/10,000 UMI for each cell. Data has been plotted as a log1p transform (applied in RStudio^21^) thereby allowing the plotting of zero values and accommodation of the broad spread of the data. All plots were prepared using ggplot2^22^ and R^23^ (https://www.R-project.org).

### Nomenclature of MHC genes

Consistent with the recommended guidelines for non-human MHC nomenclature (IPD-MHC^24,25^) mouse lemur *MHC class I* genes have the prefix “Mimu”, deriving from the genus and species <u>Mi</u>crocebus <u>mu</u>rinus. This is followed by the gene name. Pending the formal naming of the Mimu *MHC class I* genes, we retained the naming scheme of Averdam et al^11^, based upon the position of the genes within the BAC that was sequenced. We have removed the “ps” pseudogene designation as we have evidence of expression for all of the genes. The *class II* genes are named according to their homology with *MHC class II* genes in other non-human primate species.

## RESULTS

### Genomic structure of Mimu MHC class I and class II regions

Genomic sequences of the *Mimu-MHC class I* and *class II* regions were determined by Averdam et al^10,11^, who sequenced a series of seven BAC clones containing *MHC class I* or *II* genes. They found that the *Mimu-MHC class I* region located on chromosome 6 contains only *class I* pseudogenes. This region is syntenic to the human *MHC* region containing expressed *HLA class I* genes. We extracted the same regions covered by the BACs in the Mmur_3.0 genome assembly^15^ (https://www.ncbi.nlm.nih.gov/assembly/GCF_000165445.2) and compared them to the BAC sequences. The results (Fig. 1a) show good concordance between the BAC and genomic sequences, with no additional *Mimu-MHC class I* genes being identified. Additionally, no transcripts from the Tabula Microcebus Cell Atlas mapping to the homologous *class I* gene sequences of this region were obtained, confirming that this region contains only *class I* pseudogenes. We found the *MOG* gene, often considered a boundary of the *MHC class I* region. It is annotated to be 172kb from the end of mouse lemur chromosome 6, whereas its human counterpart is 29Mb from the chromosome’s terminus. The upstream genes of human chromosome 6, such as TRIM27 and the BTN gene cluster, locate to mouse lemur chromosome 14. Given the proximity of this region to the telomere in the mouse lemur, it is possible that the *class I* genes present in the *MHC* region have been inactivated by genomic instability.

**Figure 1.**
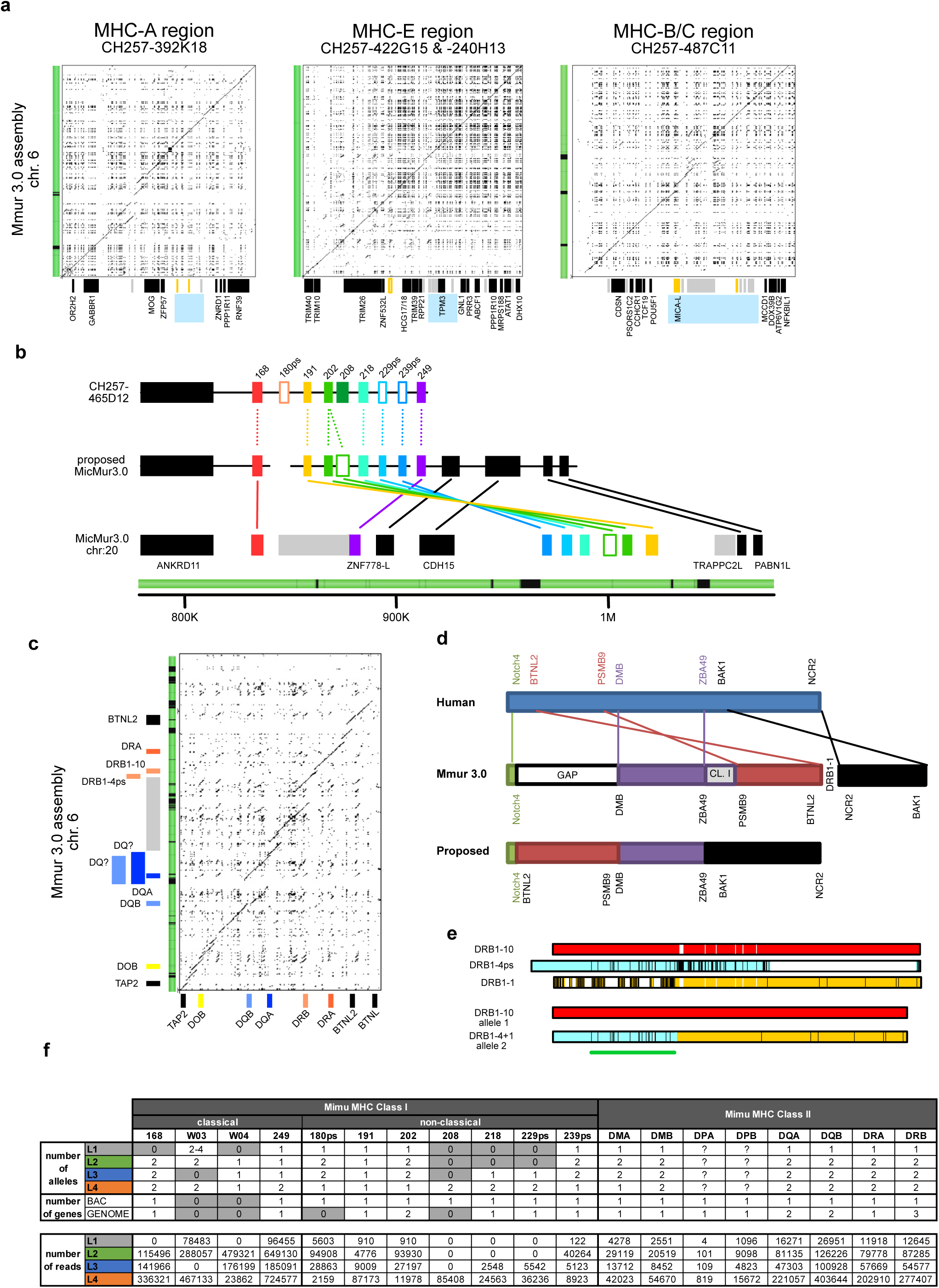
Genotyping Mimu-MHC class I and II. (a) Dotplots comparing whole genome assembly and previously sequenced BACs for the MHC class I region. The bars on the left are taken from the genome assembly: green indicates assembled regions and black indicates regions with gaps in the assembly. At the bottom of each plot, the locations of genes in the interval are indicated by boxes. Black boxes are genes predicted to be expressed, gray boxes are pseudogenes and gold boxes indicate MHC class I pseudogenes. No expressed class I genes are present in this region. Blue boxes denote regions containing expressed class I genes in humans. (b) A schematic comparing the genome assembly with the sequenced BACs that contain the predicted expressed MHC class I genes of the mouse lemur. The genes are color-coded and open boxes represent genes annotated as pseudogenes. The bar below shows regions of assembly (green) and gaps in the assembly (black). (c) Dotplot comparison of the genome assembly and previously sequenced BAC (horizontal axis). Genes are color-coded. The bar to the left of the plot shows regions of assembly (green) and gaps (black). (d) Schematic representation of the class II region of mouse lemur (upper; genome assembly), human (middle) and the proposed corrected mouse lemur organization (lower). (e) Comparison of the coding regions of the three DRB genes identified in the whole genome assembly. Vertical lines denote positions of difference in the alignment, with DRB1-10 representing the reference allele. The upper three bars are the genes annotated in the genome assembly. The lower two bars show the comparison of DRB1-10 and the composite allele obtained by combining DRB1-1 and DRB1-4. The green line indicates sequence corresponding to exon 2 encoding the polymorphic beta-1 domain. (f) Number of alleles for each locus genotyped (upper panel) and number of reads mapping to each locus (lower panel). The uncertainty in allele number for L1 is due to ambiguity in the sequencing at a single position in each of two alleles (Supp. Fig. 1). This position is in a homopolymer tract and could be a sequencing artifact or a polymorphism.

The functional *Mimu-MHC class I* genes are annotated on chromosome 20 of the genome assembly. In contrast, Averdam et al used a FISH assay to map these genes to lemur chromosome 26q^11^. The BAC sequence for this region^11^ has non-*MHC* sequences at one end of the BAC and the *Mimu-249* gene at the other end. We compared the sequenced BAC to the genome assembly (Fig. 1b). Phylogenetic comparison of the *Mimu-MHC class I* genes in the BAC with those in the genome assembly, shows that the latter lacks homologs of the *Mimu-180* and *Mimu-208* class I genes of the BAC. The genome assembly also contains a duplication of the *Mimu-202* gene (Supplemental Figure 1). This could be due either to genomic polymorphism or incorrect assembly. In the BAC, the *ANKRD11* gene is next to the *MHC* locus and located upstream of the 5’end of the first *MHC class I* gene in the cluster. Similarly, in the genome assembly, the ANKRD11 gene is followed by two closely spaced *Mimu-MHC class I* genes. Phylogenetic analysis shows these two genes are most similar to the *Mimu-168* and *Mimu-249* genes of the BAC, respectively. These genes define the boundaries of the *Mimu-MHC class I* gene cluster sequenced from the BAC. Further downstream in the genome assembly, and separated by intervening non-MHC genes, is a cluster of six *Mimu-MHC class I* genes that are homologous to five of the genes situated between *Mimu-168* and *Mimu-249* in the BAC. In the genome assembly, this gene cluster is flanked by gaps and their gene order is the inverse of that in the BAC. As these features are consistent with an assembly misalignment, we inverted the cluster and placed it between the *Mimu-168* and *Mimu-249* like genes, making it consistent with the BAC sequence. This alternative assembly brings the flanking non-MHC genes together in the order seen on human chromosome 16.

Similarly, some errors in assembly of the *Mimu-MHC class II* region have likely occurred. In addition to the many gaps in this region, the dotmatcher analysis (Fig.1c) showed several areas of possible duplication in comparison to the BAC sequence^10^. In addition, comparing gene order to the corresponding region of the human genome identified candidate regions, in which the sequence is either rearranged or misassembled (Fig. 1d). A likely contributor to the difficulty of assembling this region is the complexity of the allelic polymorphism of *Mimu-DRB*^4–7^. Genotyping mouse lemur populations has identified numerous *Mimu-DRB* alleles, with >100 alleles now deposited in GenBank^4–7^. Such diversity could have led to the errors in assembly shown in Fig. 1e. Of the three *Mimu-DRB* genes present in the genomic assembly, only *Mimu-DRB1-10* is full length. The other two genes have non-MHC sequence at either the 5’ (*DRB1-1*) or 3’ (*DRB1-4*) end. These breaks in homology both occur at the same location in intron 2. Combining the two sequences gives a complete second *Mimu-DRB* allele (Fig. 1e).

### *Assessing expression and allelic variation of individual* MHC *genes*

A reference dataset containing the known *Mimu-MHC class I* and *II* genes (Supp. Table I) was used to extract and map reads from the 10x datasets for each individual lemur (231,752 cells analyzed in total). Confirming their status as pseudogenes, no reads mapping to the *Mimu-MHC class I* genes located on chromosome 6 were obtained, neither did any reads map to *Mimu-DOA* or *Mimu-DOB*. In contrast, reads were obtained for other *Mimu-MHC* genes. In addition, reads mapping to genes identified as pseudogenes by Averdam et al^11^ were found to be expressed by individuals in our analyses. We have dropped the “ps” pseudogene designation in our discussion of these genes.

Following the initial screen, read mapping was used to assess the allelic variability. Because the *MHC class II* genes differ in sequence, ambiguous mapping was not a problem and we were able to assess allelic variability across entire genes. Although greatest read depth was found in the 600-650 bp region at the 3’ end of the cDNA sequence, as expected for the 10x data, there was sufficient read depth at the 5’ end of *MHC class II* genes to determine allelic variation. Among the four individuals there are four *MHC class II* haplotypes (*LH1, LH2, LH3* and *LH4*). Lemur L1 is homozygous for all the *Mimu-MHC class II* genes and is homozygous for haplotype *LH1*. Individuals L2 and L3 are identical and heterozygous for all *class II* genes. This was not unexpected, because L2 and L3 are descended from a common grandparent. For each of the *class II* genes, L2 and L3 shared one allele with L1, suggesting that L1, L2 and L3 share the *LH1* haplotype and in addition L2 and L3 have the *LH2* haplotype. Individual L4 is heterozygous for all *class II* genes and shares no *MHC class II* alleles with L1, L2 or L3 and is designated as having the *LH3* and *LH4* haplotypes. In the expression analysis, the count of reads was confined to the 600-650 bp 3’ end of the cDNA sequence. This provided comparison to the *MHC class I* expression levels.

Our analysis of *Mimu-MHC class I* genes was limited to the 3’ terminal 600-650 bp, because of low sequencing depth and ambiguous mapping of reads in the remainder of the sequence. Sequence homology and recombination precluded a complete phasing of SNPs across the full length of the cDNA. There was sufficient variability between the *Mimu-168, Mimu-W03* and *Mimu-W04* sequences to use them as separate references (Supp. Fig. 1), along with *Mimu-249* (identical to *Mimu-W01* and *Mimu-W02*), to assess expression of the classical *class I* genes. Some or all of these could represent allelic variants, but further genomic and population analysis will be needed to define their status. Supporting structural variation, or the variable existence of pseudogenes, reads were not obtained for all *Mimu-MHC class I* genes in all individuals (Fig. 1f). For example, no reads mapping to *Mimu-168* were found in L1, although it is well represented in the other individuals. This absence of *Mimu-168* reads in this individual is due either to absence of the gene or to its pseudogenization and consequent loss of expression.

Contrasting with the haplotypic similarities in the *MHC class II* region, analysis of the *MHC class I* region found all four individuals to be heterozygous for one or more genes, and all haplotypes to have variable gene content. As for *class II*, L1 has the least variability, being homozygous for all but one of the *MHC class I* genes. The exception is *Mimu-W03*, for which there is evidence for 2-4 alleles. The four candidate alleles vary by 1-4 nucleotides (Supp. Fig. 1c), with only one of them (*Mimu-W03-3*) being shared with another individual (L2).

L1 and L2 have *Mimu-*191 in common and lack the *Mimu-208, -218* and *-229* genes (Supp. Fig. 1). They differ at other genes, confirming that L1 and L2 have no *MHC class I* haplotype in common. Comparing L2 to L3, the identities observed for the *MHC class II* region were not observed. These individuals differ in gene content, with L3 lacking *Mimu-W03* and expression of *Mimu-218* and *-229* in contrast to L2. L2 and L3 have alleles of five genes in common (*Mimu-168, -W04, -249, -202 and -180)*. Individual L4 has no allele in common with another lemur and is the only individual that has expression of *Mimu-208* (Fig. 1f, Supp. Fig. 1). Further highlighting the difference between L4 and the other mouse lemurs are the *Mimu-191* sequences. Comparison of the BAC and genome sequences identified two allelic lineages. L4’s sequence of this region is identical to that in the BAC but differs from that of the other three lemurs by 33-34 nucleotides. L3’s sequence of this region is identical to that in the genome assembly and differs by a single nucleotide from that of L1 and L2, which are identical to each other (Supp. Fig. 1).

In comparing the BAC and genomic sequences, two of the seven genes (*Mimu-180* and *Mimu-208*) were not annotated in the genome sequence, nor were they found by sequence search. Of these, *Mimu-180* is expressed by each of the four mouse lemurs, whereas *Mimu-208* is only expressed by L4. It is possible that *Mimu-180* was missed in the genome assembly, because its location would have been at the end of the cluster bounded by gaps (Fig. 1d). The absence of *Mimu-208* from the genome assembly likely reflects a difference in gene content compared to the haplotype present in the BAC. Supporting this is the apparent linkage between alleles of *Mimu-191* and presence/absence of *Mimu-208*. The four individuals studied form two groups based on the sequence differences described for *Mimu-191* and the presence/absence of *Mimu-208*. L4 has the same sequence as that of the BAC and is the only individual having transcripts that map to *Mimu-208*. L1, L2 and L3 lack transcripts mapping to *Mimu-208* and have *Mimu-191* sequences that are either identical to the genome sequence, or differ by only one nucleotide, but differ from the L4 sequence by 33 or 34 nucleotides. In summary, the lack of expressed *Mimu-208* in these three individuals is most likely due to absence of the *Mimu-208* gene, as is seen in the genome assembly. The presence of *Mimu-208* in L4 likely represents a haplotype having different gene content. The absence of transcripts for *Mimu-218* and *Mimu-229* could be due to absence of the genes, or lack of their expression due to an inactivating mutation. More extensive genomic analysis will be necessary to determine which of these interpretations is correct.

Our initial analysis showed how using a reference set of genes, which included genes not present in an individual, led to mismapping of reads to regions with modest homology. Increasing the stringency of mapping led, however, to a loss of reads that map to polymorphic regions. To maximize read retention, while obtaining the most accurate mapping, we used reference sets specific to each lemur based on the results of our initial analysis. Reference sequences were limited to 600-650 bp in the 3’ region of each gene. Only genes that are expressed by an individual were included. These criteria minimized mismapping caused by regions of homology. The reference sequences were used to extract reads from the individual 10x datasets. Using a custom bioinformatics pipeline (see Methods) we determined the number of reads mapping to each gene (Fig. 1f). Mapping quality scores were used to exclude reads mapping ambiguously. This strategy gave conservative estimates of the level of gene expression. Fig 1f gives the total numbers of reads recovered from the 10x dataset from which doublets were excluded (215,524 cells total). Inspection of the mapping enabled us to determine the number of alleles present for each gene (Fig. 1f).

### Patterns of classical MHC class I, non-classical MHC class I, and MHC class II expression

Classical *MHC class I* genes are, by definition, expressed on the majority of cell types and at higher levels than the *non-classical class I* genes. The latter exhibit lower levels of expression and are often expressed with tissue specificity. From the numbers of reads recovered, we sorted the *Mimu-MHC class I* genes into high-expressing classical *class I genes* (*Mimu-168, -W03, -W04, -249*) and non-classical *class I genes* (*Mimu-180, -191, -202, -208, -218, -229, -239*) having lower levels of expression.

In the mouse lemur dataset, *Mimu-249* is the only classical *MHC class I* gene expressed in all individuals and by most cell types (Fig. 1f and 2a). Three additional *MHC class I* sequences, *Mimu-168, Mimu-W03* and *Mimu-W04*, are only present or expressed in some individuals. For individuals who express these genes, a majority of cells express these genes and at levels comparable to *Mimu-249*. These three sequences are likely to represent allelic variants of one or more classical *class I* genes.

Of the seven *non-classical MHC class I* reference sequences, four are present in all four individuals studied, two are present in two individuals and one is present in a single individual (Fig. 1f and 2a). These genes are expressed by a smaller fraction of the cells analyzed (1-54%, mean 23%) and at lower levels, suggesting they represent *non-classical class I* genes.

Except for *Mimu-DPA* and *Mimu-DPB*, the majority of mouse lemur MHC *class II* genes are expressed by 10-40% (mean 22%) of the cells (Fig. 2a). Expression of the *MHC class II* genes *Mimu-DPA* and *Mimu-DPB* is restricted to few cells (<1% for *DPA* and 4-9% for *DPB*).

**Figure 2.**
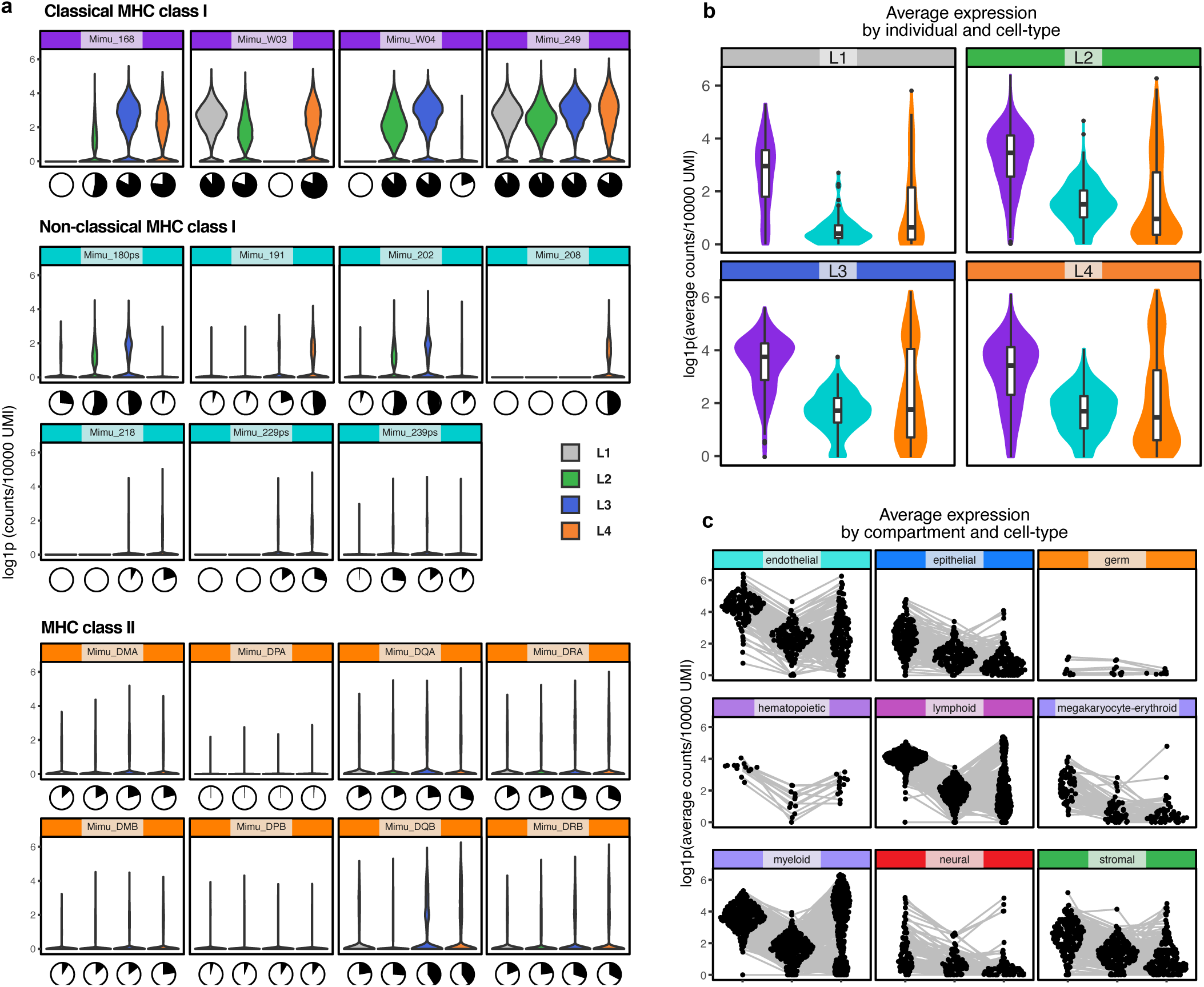
Average expression of *MHC* genes. The class I *MHC* genes are separated into classical and non-classical genes based on expression level and the frequency (percentage) of cells expressing the gene. Expression levels are compared among the four individuals (Panels a and b) and between cellular compartments (Panel c). (a) Violin plots of expression level, calculated as log1p values, for each *MHC* gene. Individuals are color-coded (gray, L1; green, L2; blue, L3; orange, L4). Genes are grouped as classical class I (purple), non-classical class I (cyan) and class II (orange). Below each violin plot, the pie chart shows the percent of cells expressing the gene shown in black. (b) Values for classical class I, non-classical class I and class II expression were summed for each cell and then used to calculate average values for individual cell types. The values are plotted as violin plots. Overall, lower values were observed for individual L1, because of the limited number of tissues examined. (c) The values plotted in panel b were then divided by cellular compartment and plotted as sina plots in panel c

### General patterns of MHC class I and class II expression

A summary dataset was created by summing the number of reads for each of the three groups (classical *class I*, non-classical *class I* and *class II*) and then normalization, as was done for the individual genes. These summed values were then averaged for groups of cells based on the individual lemur, tissue and cell-type. The summary values for each individual are shown in Fig. 2b. The patterns of expression are qualitatively similar for three of the four individuals studied. The observed differences for lemur L1 are likely due to the sampling of only three tissues from this individual. For all individuals, classical *MHC class I* genes are expressed at higher levels than non-classical *MHC class I* genes, and class II are more variable by cell type and tissue.

The summary dataset was analyzed following assignment of the cell-types into their corresponding compartments (Fig. 2c). The atlas cell types can be categorized into three distinctive types according to the pattern of MHC expression. In the first, expression is highest for classical *class I*, intermediate for non-classical *class I* and low or absent for *class II*. This hierarchy is most clearly seen in the epithelial compartment where a majority of cell-types conform to this pattern. The second most common expression pattern is characterized by high expression of both *class II and* classical *class I*, compared with intermediate expression of non-classical *class I*. This pattern is seen for large groups of cells in the endothelial and immune cell compartments with the exception of the megakaryoid/erythroid cells. The highest levels of *class II* expression are observed in dendritic cells in the myeloid compartment. This expression pattern is also characteristic of a minority of specialized cell-types in other compartments (non-myelinating Schwann cells, adipo- and osteo-CAR cells, and endothelial and AT2 cells of the lung) (The Tabula Microcebus Consortium, manuscript in preparation). Also distinguished is the third pattern comprising cells expressing low levels of *MHC class I* and *II*, a pattern most clearly seen in the germ and neural compartments.

When comparing differences in expression pattern for each cell type by tissue (Fig 3a), we observed that c*lass I* and *II* expression was lowest in tissues where the majority of cell-types were neural or germ compartment cells (brain cortex, brainstem, eye retina, hypothalamus pituitary, testes). The low levels of expression in the heart are likely due to the low quality of sequencing data from this tissue. In other tissues, the average *MHC class I* and *II* expression levels are similar. Within each tissue, the expression levels of the individual cell types varied by compartment and usually conformed to the following hierarchy: epithelial < stromal < immune types < endothelial. Classical and non-classical *MHC class I* genes conformed to this pattern. *Class II* expression was more variable, both within and between the tissues, with highest expression being seen for myeloid cells, independently of their tissue of origin. Similar expression levels were observed for subsets of endothelial cells of the kidney and lung and, to a lesser extent, the skin.

**Figure 3.**
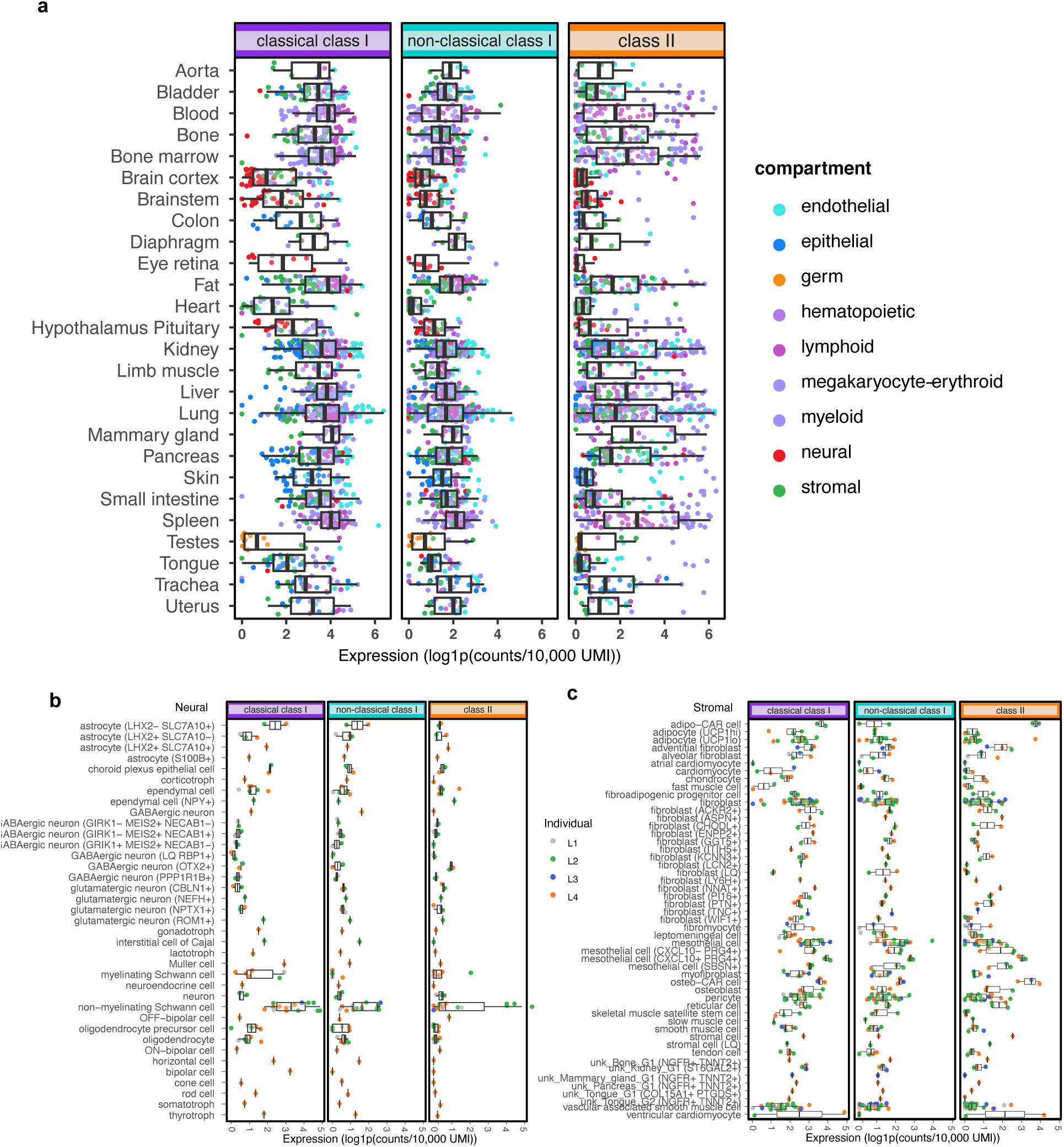
Tissue and compartment specific patterns of *MHC* expression. Each data point represents an average expression value for a cell-type, tissue or individual. (a) Comparison of expression levels among tissues. Data points are color-coded by cellular compartment. (b) Comparison of expression in cells of the neural compartment. Data points are color-coded by individual. (c) Comparison of expression in cells of the stromal compartment. Data points are color-coded by individual.

Some neural compartment cell-types showed increased expression of *MHC class I* and *II* (Fig. 3b). The non-myelinating Schwann cells have higher expression of classical MHC *class I* compared to other cell-types in the same compartment, including myelinating Schwann cells, neurons, and astrocytes. This was seen for all individuals, with L2 and L3 demonstrating the highest expression. These two lemurs also had higher levels of non-classical *class I*, than L1 and L4. Uniquely, L2 had elevated *MHC class II* expression in non-myelinating Schwann cells of the periphery (fat, pancreas, kidney and small intestine) but not in cells of the brainstem or cortex. Elevated *MHC class II* expression by non-myelinating Schwann cells has previously been observed in inflammatory responses^26^, and is consistent with L2’s known infections (cystitis and pneumonia) and overall inflammation due to uterine cancer^27^.

Most stromal cells have modest levels of *MHC class I* and low levels of *class II*. Notable exceptions are the adipo- and osteo-CAR cells which expressed high levels of *MHC class II*, consistent with them being central niche cells that are necessary for the proliferation and maintenance of hematopoietic stem cells, as well as the proliferation of lymphoid and erythroid precursors^28,29^. Also exhibiting elevated *MHC class II* expression were the PRG4+ mesothelial cells of fat and pancreas, and in particular, those expressing the chemokine ligand CXCL10 had higher *class II* expression than those lacking CXCL10.

### MHC expression along the developmental trajectory of myeloid and erythroid cells

We used the pseudo-time developmental trajectory, described in The Tabula Microcebus Consortium, et al^14^, to assess differences in *MHC* expression in the myeloid and erythroid cells of bone and bone marrow (Fig. 4). Shown here (Fig 4) are data from three individuals, for which trajectories were constructed. Overall, these individuals exhibit similar patterns of gene expression. Classical class I expression is highest in hematopoietic precursor cells and lowest in cells of the erythroid lineage. Class II expression is highest in monocytes and macrophages, and virtually absent from neutrophils and erythroid cells. These data are consistent with observed patterns of MHC expression in humans and mice^13,30^.

**Figure 4.**
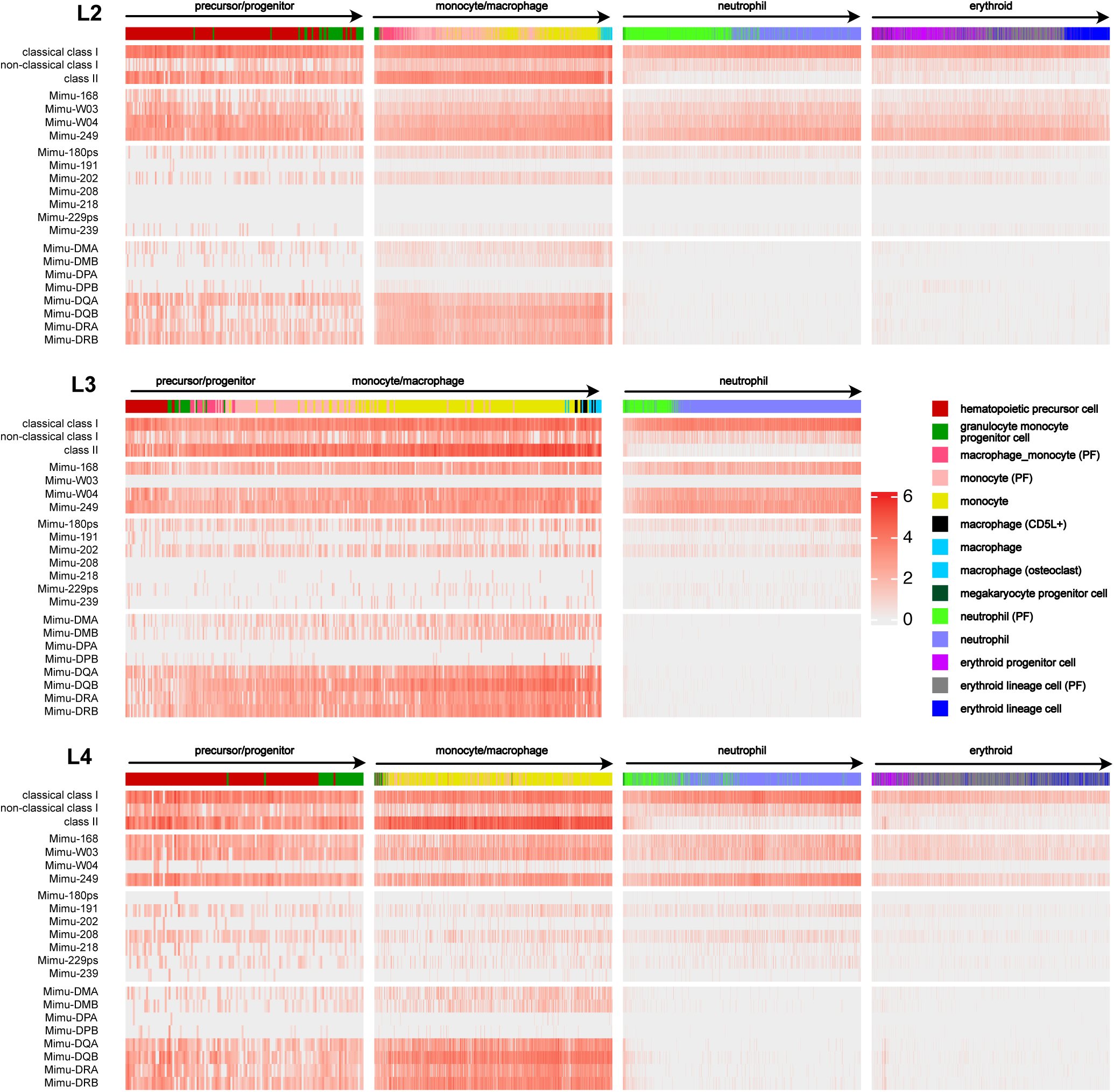
*MHC* expression in myeloid and erythroid cells from bone and bone marrow. The heat maps show the expression levels of the *MHC* genes in cells that had been placed on a trajectory^14^. In all plots the direction of the trajectory is shown by arrows and the major cell type in each trajectory is indicated. Each individual plot represents a branch of the trajectory determined within the individual. Above each heat map the cell-type is indicated by the colored bar. The upper three lines of each panel show the summed values for the expression of classical class I, non-classical class I or class II. Below are the individual values for each gene in those three groups. As noted previously for the trajectory analysis, dendritic cells did not form part of the continuum and were excluded from that analysis.

The individual lemurs had distinctive patterns of gene expression. In L2, *Mimu-168* and *Mimu-W03* were expressed at similarly low levels, whereas *Mimu-W04* and *Mimu-249* were expressed at higher levels. This suggests that *Mimu-168* and *Mimu-W03* could be alleles of the same gene, whereas *Mimu-W04* and *Mimu-249* represent different genes, each present as two copies. In L3 there was no expression of *Mimu-W03* and in L4, *Mimu-W04* has low expression resembling that of a non-classical *MHC* class I gene. The overall expression of *non-classical MHC class I* is similar for the three lemurs. However, in L2 and L3 it was primarily due to *Mimu-180* and *Mimu-202* expression, whereas in L4 it was due to *Mimu-191* and *Mimu-208* expression. Patterns of MHC *class II* expression were similar for L2, L3 and L4, comprising low expression of *Mimu-DP*, modest expression of *Mimu-DM* and high expression of *Mimu-DR* and *-DQ*.

### MHC expression along the spatial gradient of the kidney nephron epithelium and endothelium

The epithelial cells of the kidney nephron form a continuum in overall gene expression that resembles their spatial gradients^14^ (data present for L2, L3, L4) (Fig. 5). Along the nephron cells, the highest levels of MHC class I and II expression are seen in the descending limb cells of the loop of Henle. In contrast, the cells of the proximal convoluted and proximal straight tubules have the least expression. This pattern was consistent among the three individuals. Similarly, highest expression of *MHC* class II occurs in cells of the thin descending loop of Henle. The vasa recta endothelial cells of the kidney also form a molecular continuum that resembles their spatial gradients from the descending (arteriole) to the ascending (venous) end. A subtle gradient of class II expression was observed with expression increasing from cells of the ascending limb to those of the descending limb. The overall levels of expression were greater in the vasa recta than in cells of the nephron epithelium, consistent with the levels observed on comparing the endothelial and epithelial compartments.

**Figure 5.**
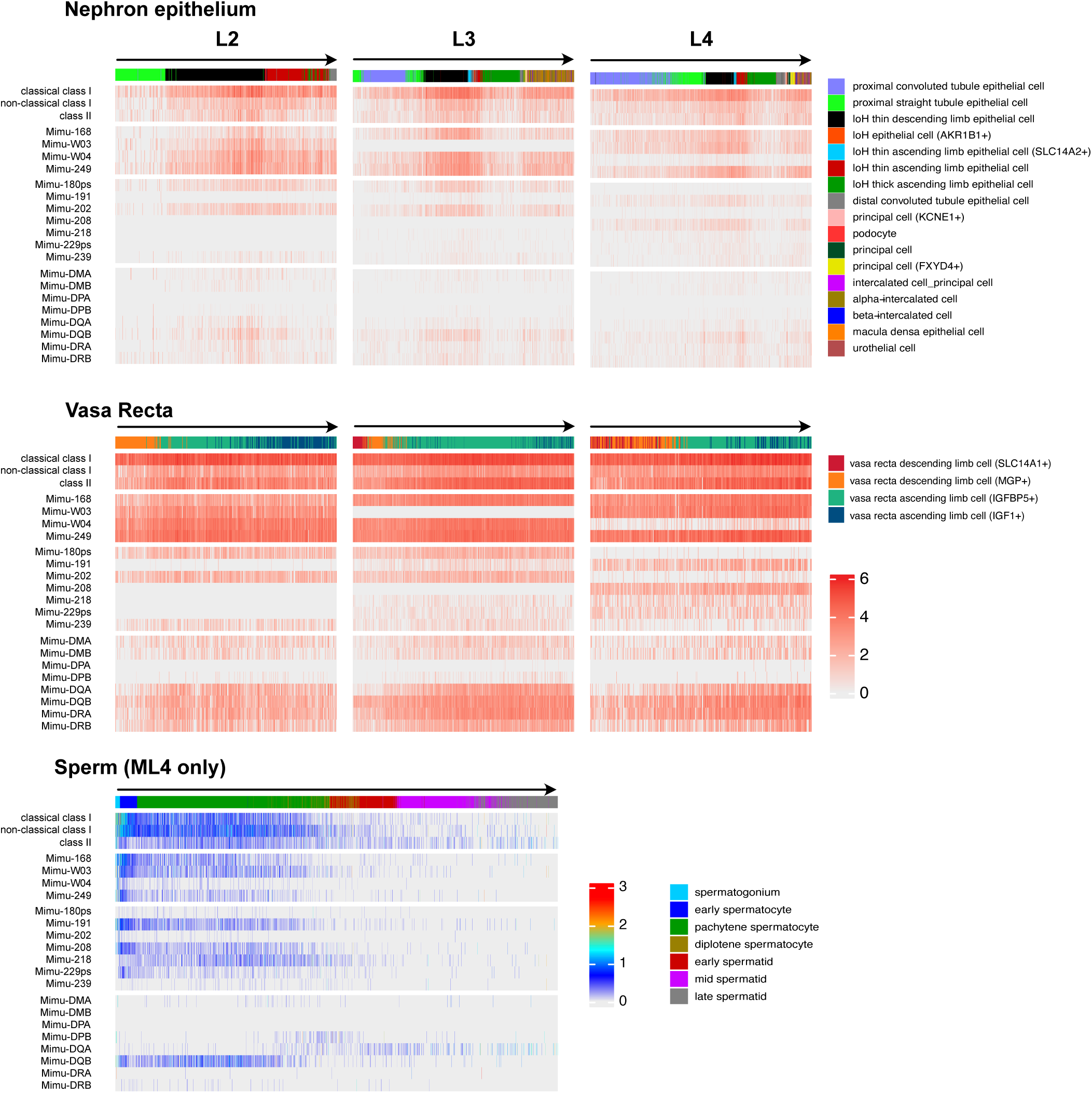
*MHC* expression in epithelial cells of the kidney and germ cells of the testis. The heat maps show the expression levels of the *MHC* genes in cells that had been placed on a trajectory^14^. In all plots the direction of the trajectory is shown by the arrows. Above each heatmap the cell-type is indicated by the colored bar. The top three lines of each panel show the summed values for classical class I, non-classical class I and class II expression. Below are the individual values for each gene in those three groups. The top group of heatmaps is for nephron epithelial cells, the middle group is vasa recta. Shown at the bottom is the heat map based on the sperm cell trajectory. The alternate color ramp for the sperm trajectory highlights the overall lower levels of expression in the sperm cells.

### MHC expression along the development trajectory of the spermatogenesis

Expression of MHC molecules during spermatogenesis has remained a topic of debate. Various studies have reported low levels of cell surface expression, no expression or expression of mRNA with no detectable cell-surface expression^31–36^. The trajectory described for the sperm cells of L4 followed their developmental pathway starting from spermatogonia and maturing into late spermatids^14^. Mapping *MHC* expression on this trajectory showed that although the levels of expression were much lower than seen for other cell compartments, specific patterns of expression could be discerned. All *classical* and most *non-classical class I* genes were most highly expressed during early development (in spermatagonia and early spermatocytes), but then declined through the pachytene spermatocytes, becoming virtually absent in later development. Surprisingly, in the developing sperm we observed expression of classical and non-classical MHC class I at similar levels, contrasting with the lower level of non-classical class I expression by most other cell types. However, individual non-classical class I genes have distinctive patterns of cellular expression. *Mimu-191* and *Mimu-208* have highest expression in early development, followed by a decline in pachytene spermatocytes and eventual loss of expression (Fig. 5). *Mimu-229* is present on spermatagonia and early spermatocytes but absent from subsequent developmental stages. *Mimu-218* is preferentially expressed by pachytene spermatocytes.

Throughout the trajectory of spermatocyte development there is a low, but detectable, level of *MHC class II* expression. As for the individual non-classical class I genes, examining the expression of individual class II genes showed clear differences, particularly between Mimu-DQA and DQB. *Mimu-DQB* is expressed by early spermatocytes and then declines in the pachytene spermatocytes and is associated with little expression of *Mimu-DQA*. Surprisingly, beginning with the early spermatids and continuing in the mid spermatids, there is increasing expression of *Mimu-DQA*, with little expression of *DQB*. That these data were obtained from analysis of one geriatric individual, means that future study of younger, healthier lemurs is needed to determine if the unexpected discordance between DQA and DQB expression represents physiological mouse lemur spermatogenesis.

## DISCUSSION

We developed a bioinformatics pipeline that measures expression levels of mouse lemur *MHC class I* and *II* genes from 10x single cell RNA sequencing data. Our comparison of the genome assembly, previously sequenced BACs and our expression data confirm that the mouse lemur *MHC* region syntenic to that of human and mouse contains no expressed *MHC class I* genes. Instead, the expressed mouse lemur *MHC class I* genes all cluster on a chromosome separate from the *MHC* region. By contrast, the mouse lemur *MHC* region does contain expressed *MHC class II* genes. Comparing expression of individual genes, among the four individuals (L1, L2, L3 and L4) included in Tabula Microcebus, revealed differences that are due either to different MHC gene content or differential expression of the individual genes.

Four allelic *MHC class II* haplotypes (*LH1, LH2, LH3* and *LH4*) were identified in the four mouse lemurs studied (L1, L2, L3 and L4). L1 is homozygous for the *LH1* haplotype and shares that haplotype with individuals L2 and L3. Individuals L2 and L3 have identical *MHC class II* types. They have the *LH1* haplotype in common with mouse lemur L1 the *LH2* haplotype. Individual L4 has two *MHC class II* haplotypes *LH3 and LH4*, neither of which is present in individuals L1, L2 or L3. The mouse lemur *MHC class I* genes are more variable than the *MHC class II* genes, both in gene content and in allelic polymorphism. Although various genes and alleles are held in common, there is no sharing of *MHC* haplotypes. As for *MHC class II*, L4 shares no *MHC class I* allele with L1, L2 or L3.

The *MHC* expression data from the Tabula Microcebus were used to determine average expression levels for the various cell-types. For the four mouse lemurs, a variety of cell types were compared for their levels of expression. In general, a concordance in the levels of expression for each cell type was observed between individuals. Notable exceptions were the non-myelinating Schwann cells, where comparing the expression data for the four mouse lemurs, we found an elevation of *MHC class II* expression in one individual. This difference could have been due to an ongoing disease process in this individual lemur^27^, indicating the possibility of determining aberrant expression levels that correlate with ongoing disease.

The observed patterns of expression largely agree with known expression patterns for human and mouse. *MHC class I* is expressed across all cell-types with some variation in levels of expression (Fig. 2a). *Class II* expression is restricted to a subset of cell-types, primarily cells of immune compartments. A few exceptional cell-types were detected in other compartments that expressed *class II* at elevated levels. These included adipo- and osteo-CAR cells, PRG4+ mesothelial cells, non-myelinating Schwann cells (in a single individual), AT2 cells of the lung and endothelial cells of the capillaries, arteries and veins of the lung and kidney. None of these were unexpected, although basal *class II* expression in endothelial cells is absent in murids^28,37–40^.

By combining the expression data with previous trajectory analysis, we detected gradients of *MHC class I* and *II* expression along developmental and spatial pathways. In the nephron epithelium and vasa recta, gradients of expression were uncovered and found to be similar in all individuals studied. Previous analyses have been primarily immunohistochemical or expression analyses of renal cell carcinomas with a focus on *class II* expression^40,41^. None of these studies report the gradient of expression that we observe. The pathway of sperm development was studied in one of the lemurs. Mapping *MHC* expression to this developmental pathway revealed unexpected patterns of expression. There was separate expression of *Mimu-DQB* and *Mimu-DQA* that corresponded to specific developmental stages. Ordinarily there is coordinate expression of these genes that encode the alpha and beta chains of Mimu-DQ. As only one geriatric individual was studied, further analysis will be necessary to determine if this is a common pattern of expression or whether it was specific to this individual.

We have presented an iterative method for determining expression levels for the highly homologous and highly polymorphic *MHC class I* and *class II* genes of the mouse lemur from short read single cell RNA-sequencing data. This pipeline could be used to study *MHC* expression in other mouse lemurs as well as being expanded for use with other species. Here we have shown several representative analyses highlighting the utility of the cell atlas for comparing expression profiles between individuals, between cell-types, between tissues and between genes. The depth and breadth of the data available make detailed comparison of individuals and cell-types possible and will form the basis for future investigations of the functional consequences of the observed variations.

