## Supplemental Figure 1 for "Organism-wide mapping of MHC class I and II expression in mouse lemur cells and tissues"

**Supplemental Figure 1.** Comparison of the 650 bp tail segment for the expressed *MHC class I* genes of mouse lemur. Sequences from the genes on chromosome 6 were not included in this alignment. (A) Shown are positions of variation in an alignment of all *MHC class I* sequences present in the genome assembly, sequenced BAC (ref) and present in GenBank (W01-04, ref). The genome sequences are indicated by accession number. Note that XM_012737263 is only 200 bp in length as this was the sequence present in the genome annotation. Sequences are divided into three groups. The top group contains all of the *classical class I* reference sequences, the second group contains all of the *non-classical class I* reference sequences and the third group contains additional sequences that were identical to one of the reference sequences, so not included in further analysis. Two of the sequences had segments that are highly divergent, as indicated. Pale gray shading indicates regions for which no sequence was present. (B) Pairwise analysis showing numbers of differences between genes. Identical sequences (0 differences) are indicated in gold. It should be noted that differences exist outside of the 650 bp region included in this analysis. (C) Allelic variants of *classical class I* genes obtained in this study. Only variable positions are shown. The individual(s) from which each allele was recovered is shown by colored boxes to the left of the sequences. (D) Allelic variants of non-*classical class I* genes obtained in this study. Only variable positions are shown. The individual(s) in which each allele was recovered is shown by colored boxes to the left of the sequences.

A

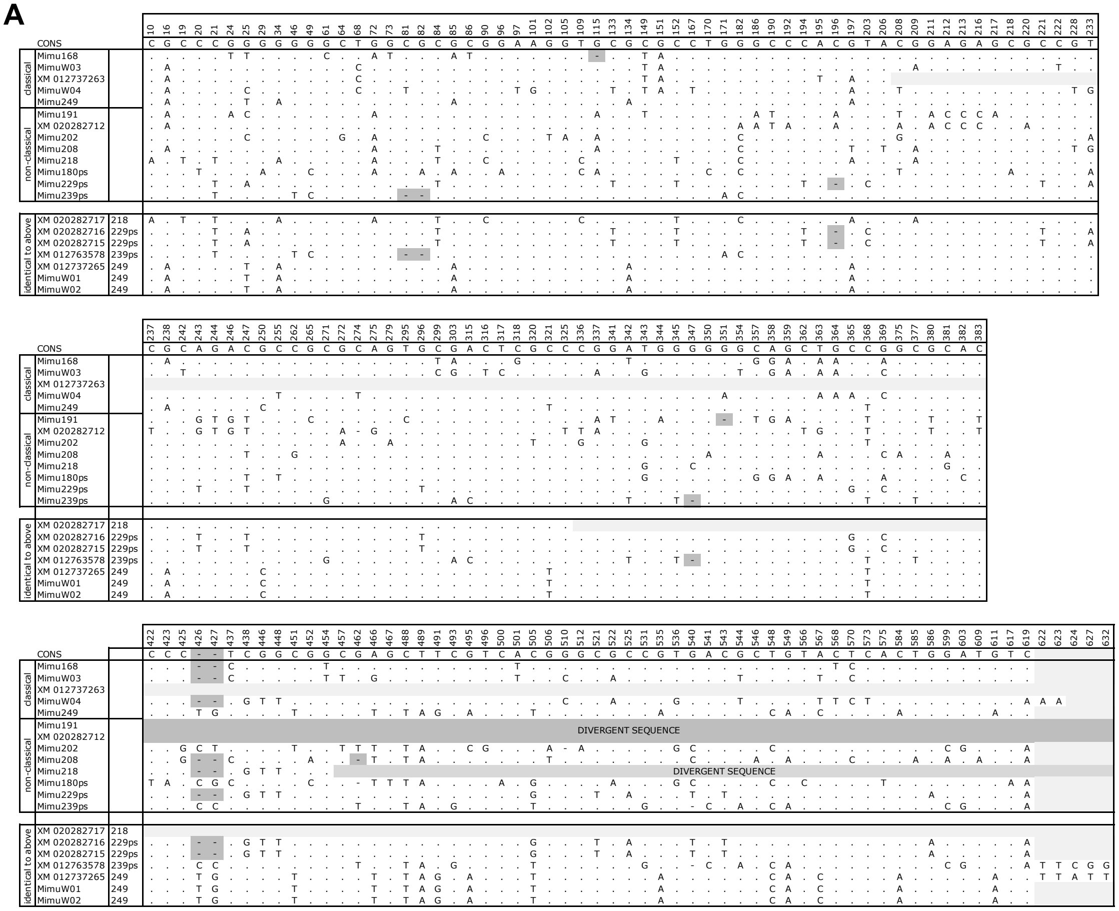


B

| number of differences; note XM_012737263 is only 200 bp in length | Mimu168 | MimuW03 | XM_012737263 | MimuW04 | Mimu249 | Mimu191 | XM_020282712 | Mimu202 | Mimu208 | Mimu218 | Mimu180ps | Mimu229ps | Mimu239ps | XM_020282717 | XM_020282716 | XM_020282715 | XM_012763578 | XM_012737265 | MimuW01 | MimuW02 |
|---|---|---|---|---|---|---|---|---|---|---|---|---|---|---|---|---|---|---|---|---|
| Mimu168 | | | | | | | | | | | | | | | | | | | | |
| MimuW03 | 44 | | | | | | | | | | | | | | | | | | | |
| XM_012737263 | 20 | 2 | | | | | | | | | | | | | | | | | | |
| MimuW04 | 57 | 36 | 7 | | | | | | | | | | | | | | | | | |
| Mimu249 | 66 | 57 | 8 | 61 | | | | | | | | | | | | | | | | |
| Mimu191 | 206 | 193 | 11 | 202 | 187 | | | | | | | | | | | | | | | |
| XM_020282712 | 165 | 144 | 22 | 152 | 134 | 49 | | | | | | | | | | | | | | |
| Mimu202 | 81 | 68 | 14 | 71 | 47 | 188 | 136 | | | | | | | | | | | | | |
| Mimu208 | 63 | 49 | 12 | 53 | 58 | 199 | 148 | 60 | | | | | | | | | | | | |
| Mimu218 | 141 | 122 | 15 | 124 | 128 | 238 | 151 | 131 | 122 | | | | | | | | | | | |
| Mimu180ps | 74 | 66 | 17 | 72 | 55 | 195 | 150 | 53 | 61 | 138 | | | | | | | | | | |
| Mimu229ps | 62 | 51 | 14 | 44 | 58 | 197 | 148 | 67 | 51 | 114 | 69 | | | | | | | | | |
| Mimu239ps | 80 | 74 | 17 | 77 | 50 | 199 | 146 | 53 | 70 | 134 | 64 | 68 | | | | | | | | |
| XM_020282717 | 32 | 19 | 15 | 26 | 17 | 32 | 44 | 19 | 17 | 0 | 22 | 20 | 22 | | | | | | | |
| XM_020282716 | 62 | 51 | 14 | 44 | 58 | 198 | 148 | 67 | 51 | 114 | 69 | 0 | 68 | 20 | | | | | | |
| XM_020282715 | 62 | 51 | 14 | 44 | 58 | 198 | 148 | 67 | 51 | 114 | 69 | 0 | 68 | 20 | 0 | | | | | |
| XM_012763578 | 80 | 74 | 17 | 79 | 50 | 202 | 146 | 53 | 70 | 138 | 64 | 68 | 0 | 22 | 68 | 68 | | | | |
| XM_012737265 | 66 | 57 | 8 | 63 | 0 | 191 | 134 | 47 | 58 | 131 | 55 | 58 | 50 | 17 | 58 | 58 | 51 | | | |
| MimuW01 | 66 | 57 | 8 | 61 | 0 | 187 | 134 | 47 | 58 | 128 | 55 | 58 | 50 | 17 | 58 | 58 | 50 | 0 | | |
| MimuW02 | 66 | 57 | 8 | 61 | 0 | 187 | 134 | 47 | 58 | 128 | 55 | 58 | 50 | 17 | 58 | 58 | 50 | 0 | 0 | |

C

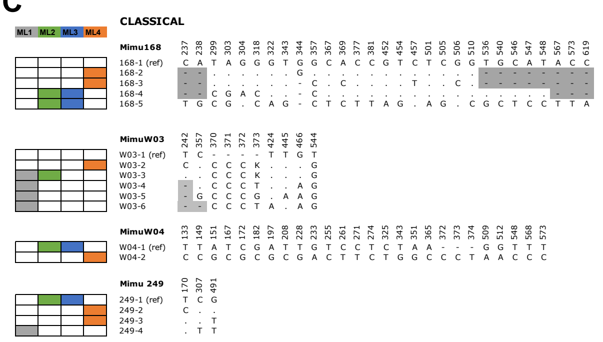


D

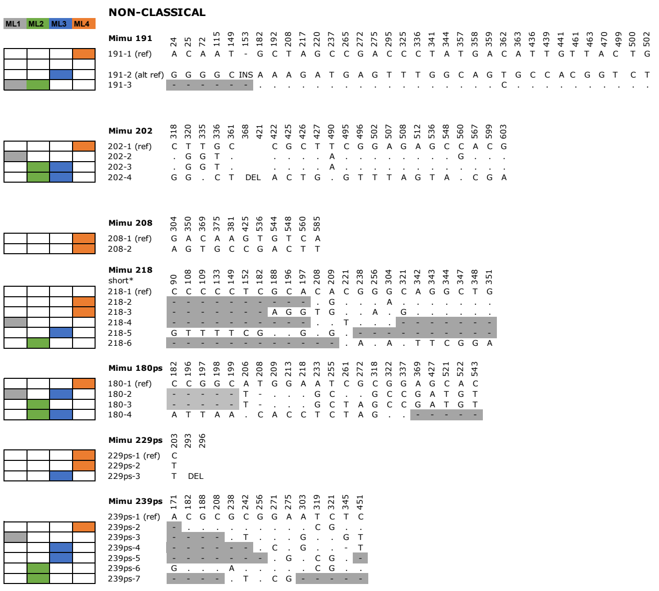
