## Supplemental Table 1 for "Organism-wide mapping of MHC class I and II expression in mouse lemur cells and tissues"

**Supplemental Table 1.** Sequences used for bowtie2 index for read harvesting.

| Gene | Accession | Notes |
|---|---|---|
| chr.6 MHC class I from Mmur_3.0 assembly | | |
| pseudo | LOC105855356 | in region syntenic to HLA-A,-F,-G,-H |
| pseudo | LOC105855357 | in region syntenic to HLA-A,-F,-G,-H |
| pseudo | XM_020287333.1 | LOC105858107; in region syntenic to HLA-B |
| chr.20/26MHC class I from Mmur_3.0 assembly | | |
| Mimu-168 | FP236833 | from sequenced BAC |
| Mimu-168 | LOC105855949 | |
| Mimu-180 | FP236833 | from sequenced BAC |
| Mimu-191 | FP236833 | from sequenced BAC |
| Mimu-191 | LOC105870762 | |
| Mimu-202 | FP236833 | from sequenced BAC |
| Mimu-202 | LOC105870767 | |
| Mimu-202 | LOC105870769 | |
| Mimu-208 | FP236833 | from sequenced BAC |
| Mimu-218 | FP236833 | from sequenced BAC |
| Mimu-218 | LOC105810165 | |
| Mimu-229 | FP236833 | from sequenced BAC |
| Mimu-229 | LOC105870764 | |
| Mimu-239 | FP236833 | from sequenced BAC |
| Mimu-239 | LOC105870766 | |
| Mimu-249 | FP236833 | from sequenced BAC |
| Mimu-249 | LOC105855951 | |
| Mimu-W01 | AJ297588 | cDNA sequence; likely allele of Mimu-249 |
| Mimu-W02 | AJ297589 | cDNA sequence; likely allele of Mimu-249 |
| Mimu-W03 | AJ297590 | cDNA sequence; likely allele or duplication of Mimu-168 |
| Mimu-W04 | AJ302085 | cDNA sequence; likely allele or duplication of Mimu-168 |
| chr.6 MHC class II from Mmur_3.0 assembly | | |
| DRB | XM_012765726 | DRB1-1; partial likely second allele; (LOC105872012) |
| DRB | XR_002223954 | DRB1-4; partial likely second allele; (LOC109730834) |
| DRB | XM_020286955 | DRB1-10; (LOC105876782) |
| DMA | XM_012791371 | (LOC105886004) |
| DOA | XM_012791383 | (LOC105886015) |
| DPA | XM_020286938 | (LOC105886009) |
| DQA | XM_020286957 | DQA haplotype D (LOC105869753) |
| DQA | XM_020286956 | (LOC105869752) |
| DRA | XM_012774641 | (LOC105876793) |
| DMB | XM_012791379 | (LOC105886010) |
| DOB | XM_012791391 | (LOC105886027) |
| DPB | XM_012791375 | (LOC105886005) |
| DQB | XM_012761793 | (LOC105869751) |

| | | |
|---|---|---|
| DQB | XM_020286954 | (LOC105886026) |
| MHC class II cDNA (mostly exon 2 only) | | |
| DRB | LN610539 | MibeDRB*001 |
| | LN610542 | MibeDRB*004 |
| | HE801960 | MimuDRB*066 |
| | HE801959 | MimuDRB*065 |
| | LN610553 | MibeDRB*015 |
| | HE801958 | MimuDRB*064 |
| | HE801963 | MimuDRB*069 |
| | LN610548 | MibeDRB*010 |
| | AJ555836 | MimuDRB*7 |
| | LN610550 | MibeDRB*012 |
| | AJ555839 | MimuDRB*10 |
| | HE801954 | MimuDRB*060 |
| | AJ555835 | MimuDRB*4 |
| | LN610543 | MibeDRB*005 |
| | LN610554 | MibeDRB*016 |
| | LN610544 | MibeDRB*006 |
| | LN610551 | MibeDRB*013 |
| | LN610547 | MibeDRB*009 |
| | AJ431269 | MimuDRB*5 |
| | AJ431270 | MimuDRB*6 |
| | LN610549 | MibeDRB*011 |
| | LN610541 | MibeDRB*003 |
| | AJ431266 | MimuDRB*1 |
| | HE801962 | MimuDRB*068 |
| | HE801956 | MimuDRB*062 |
| | HE801964 | MimuDRB*070 |
| | HE801955 | MimuDRB*061 |
| | HE801961 | MimuDRB*067 |
| | AJ431267 | MimuDRB*2 |
| | AJ431268 | MimuDRB*3 |
| | HE801957 | MimuDRB*063 |
| | AB078195 | MiruDRB*Wd01 |
| | AB078196 | MiruDRB*Wc02 |
| | AB078295 | MiruDRB*Wc03 |
| | AB078297 | MiruDRB*Wc04 |
| | AB078299 | MiruDRB*Wb01 |
| | AB078290 | MimyDRB*Wa03 |
| | AB078291 | MimyDRB*Wa04 |
| | AB078203 | MimyDRB*Wd01 |
| | AB078276 | MiruDRB*Wa01 |

| | | |
|---|---|---|
| DRB cont. | AB078222 | MimyDRB*Wa01 |
| | AB078256 | MiruDRB*Wg01 |
| | AJ555838 | MimuDRB*9 |
| | LN610545 | MibeDRB*007 |
| | LN610552 | MibeDRB*014 |
| | LN610546 | MibeDRB*008 |
| | AJ555837 | MimuDRB*8 |
| | AJ830741 | MimuDRB*16 |
| | AB078284 | MiruDRB*Wa01 |
| | LN610540 | MibeDRB*002 |
| | AJ555840 | MimuDRB*13 |
| | AJ555841 | MimuDRB*14 |
| | AJ830740 | MimuDRB*12 |
| DQA | HQ222964 | MimTDQA*013 |
| | HQ222957 | MimTDQA*006 |
| | HQ222959 | MimTDQA*008 |
| | HQ222958 | MimTDQA*007 |
| | HQ222956 | MimTDQA*005 |
| | HQ222961 | MimTDQA*010 |
| | HQ222962 | MimTDQA*011 |
| | HQ222960 | MimTDQA*009 |
| | HQ222955 | MimTDQA*004 |
| | HQ222963 | MimTDQA*012 |
| | HQ222938 | MimTDQA*003 |
| | HQ222936 | MimTDQA*001 |
| | HQ222937 | MimTDQA*002 |
| DRA | HQ222945 | MimTDRA*003 |
| | HQ222944 | MimTDRA*002 |
| | HQ222943 | MimTDRA*001 |
| DQB | HQ222950 | MimTDQB*006 |
| | HQ222951 | MimTDQB*007 |
| | HQ222952 | MimTDQB*008 |
| | HQ222954 | MimTDQB*010 |
| | HQ222947 | MimTDQB*003 |
| | HQ222946 | MimTDQB*002 |
| | HQ222949 | MimTDQB*005 |
| | HQ222948 | MimTDQB*004 |
| | HQ222953 | MimTDQB*009 |
| | HQ222939 | MimTDQB*001 |
